# PP1 inhibitor-2 promotes PP1γ positive regulation of synaptic transmission

**DOI:** 10.1101/2022.02.10.480004

**Authors:** Karl Foley, Haider Altimimi, Hailong Hou, Yu Zhang, Cody McKee, Makaía M. Papasergi-Scott, Hongtian Yang, Abigail Mayer, Nancy Ward, David M. MacLean, Angus C. Nairn, David Stellwagen, Houhui Xia

## Abstract

Inhibitor-2 (I-2) is a prototypic inhibitor of protein phosphatase-1 (PP1), a major serine-threonine phosphatase that regulates synaptic plasticity and learning and memory. Although I-2 is a potent inhibitor of PP1 in vitro, our previous work has elucidated that, in vivo, I-2 may act as a positive regulator of PP1. Here we show that I-2 and PP1γ, but not PP1α, positively regulate synaptic transmission in hippocampal neurons. Moreover, we demonstrated that I-2 enhances PP1γ interaction with its major synaptic scaffold, neurabin, by Förster resonance energy transfer (FRET)/Fluorescence lifetime imaging microscopy (FLIM) studies, while having a limited effect on PP1 autoinhibitory phosphorylation. Furthermore, our study indicates that the effect of I-2 on PP1 activity in vivo is dictated by I-2 threonine-72 phosphorylation. Our work thus demonstrates a molecular mechanism by which I-2 positively regulates PP1 function in synaptic transmission.

## Introduction

Inhibitor-2 (I-2) is a prototypic inhibitor of protein phosphatase-1 (PP1), a major serine-threonine phosphatase which plays a critical role in synaptic functions [1]. The mechanism of inhibition of PP1 by I-2 has been extensively studied in vitro for decades since its purification in 1976 [2, 3]. Based on the crystal structure of the PP1–I-2 complex [4] and other biochemical studies, PP1 binds tightly to I-2 in a 1:1 stoichiometry, and an α-helix of I-2 (130-169) covers the active site of PP1, thereby inhibiting catalytic activity.

PP1 is inactive within the in vitro PP1–I-2 complex, but can be quickly (<1 minute) activated when I-2 is phosphorylated at threonine 72 (pT72) by GSK3β [2], MAPK or CDK5 [5], presumably moving the I-2 α-helix away from the active site of PP1. However, I-2 pT72 acts as an intramolecular substrate for active PP1, and therefore must undergo dephosphorylation before PP1 is active towards other substrates. While PP1 is active, dephosphorylated I-2 T72 will lead to another slow (∼30 minutes) conformational change such that the α-helix of I-2 (residues 130-169) swings back into the active site, eventually leading to full inhibition of phosphatase activity [2], completing the PP1 activation-inhibition cycle. Mutation of I-2 at T72 (T72A) blocks the activation step in the cycle and should be a constitutive PP1 inhibitor [6, 7].

While in vitro studies predominate, studies of I-2 function in intact cells and model organisms suggest more complex I-2 regulation and function [8, 9]. Increasing evidence suggests that I-2 can function as a positive regulator of PP1 in addition to its role in PP1 inhibition [3, 10]. For example, we found that I-2 knockdown, but not I-1 knockdown, in primary cortical neurons decreases PP1 activity based on increased PP1 inhibitory phosphorylation at T320 [11]. Further, PP1 activity is required for long-term depression (LTD), but chemical LTD is defective in I-2 knockdown neurons [11]. Similarly, PP1 constrains learning and memory and acts as a molecule of forgetfulness [12], but global knockout (KO) of I-2 in mice, or knockdown of I-2 in rats, decreased memory formation as assayed by novel object recognition, contextual fear conditioning, and Morris water maze [13].

While we have found that I-2 plays an important role in LTD [11], synaptic downscaling [14], and memory formation [13], whether I-2 plays a role in regulating synaptic transmission has not been examined. Additionally, the mechanism by which I-2 can promote PP1 function in the central nervous system is not clear, but, like other PP1 regulatory proteins, likely involves changing PP1 interaction with other proteins. Previous research has shown that I-2 can form a heterotrimeric complex with neurabin and PP1 [15, 16], presenting an enticing model by which I-2 could regulate synaptic PP1.

Neurabin is a major PP1 regulatory protein that binds F-actin and targets PP1 to synaptic spines. Neurabin binds to PP1 via its RvXF^460^ motif as well as adjacent disordered regions [17]. Mutating F460 in the RvXF^460^ motif leads to robust and significant decrease of neurabin-PP1 binding [17, 18], as well as a decrease in synaptic transmission, suggesting that PP1 bound to neurabin promotes synaptic transmission [19]. Neurabin binds PP1γ preferentially, PP1α to a lesser extent, and PP1β minimally [20, 21], suggesting PP1γ is most likely the PP1 isoform that promotes synaptic transmission. However, no direct test has validated this. Moreover, I-2, PP1, and neurabin all localize to excitatory synapses [22, 23], suggesting that they could act together in regulating synaptic transmission.

In this study, we employed over-expression of I-2 in combination with PP1α, PP1γ, and I-2 KO studies and found that I-2 and PP1γ both promote basal synaptic transmission. By introducing a phospho-null mutation at T72 in I-2, thereby disrupting the activation-inhibition cycle of PP1–I-2, we abolished the positive effect of I-2 on PP1 activity. Lastly, we find that I-2 promotes PP1γ targeting to neurabin, a critical synaptic PP1γ scaffolding protein for synaptic transmission. Our current work elucidates an important function of I-2 in promoting synaptic transmission and provides a mechanism of how I-2 positively regulates PP1 function.

## Results and Discussion

In order to study the potential role of I-2 in regulating basal synaptic transmission, we expressed CFP (control), CFP-tagged wild-type (WT) I-2, or CFP-tagged I-2^T72A^ in primary cortical neurons and performed whole-cell recordings 3 days after transfection (**Fig. 1A**). We found that mEPSC amplitude, but not frequency, was robustly increased in CFP-I-2-expressing neurons compared to control neurons (**Fig. 1B-D**). Interestingly, neither mEPSC amplitude nor mEPSC frequency showed significant change in CFP-I-2^T72A^-expressing neurons (**Fig. 1B-D**). Our data thus suggest that I-2 promotes synaptic transmission, but this effect depends on the T72-mediated activation-inhibition cycle. While the mean mEPSC frequency was not significantly different overall, a subset of CFP-I-2^WT^-expressing neurons showed a robust increase in mEPSC frequency, though this potential effect was not further pursued in this study.

**Figure 1:**
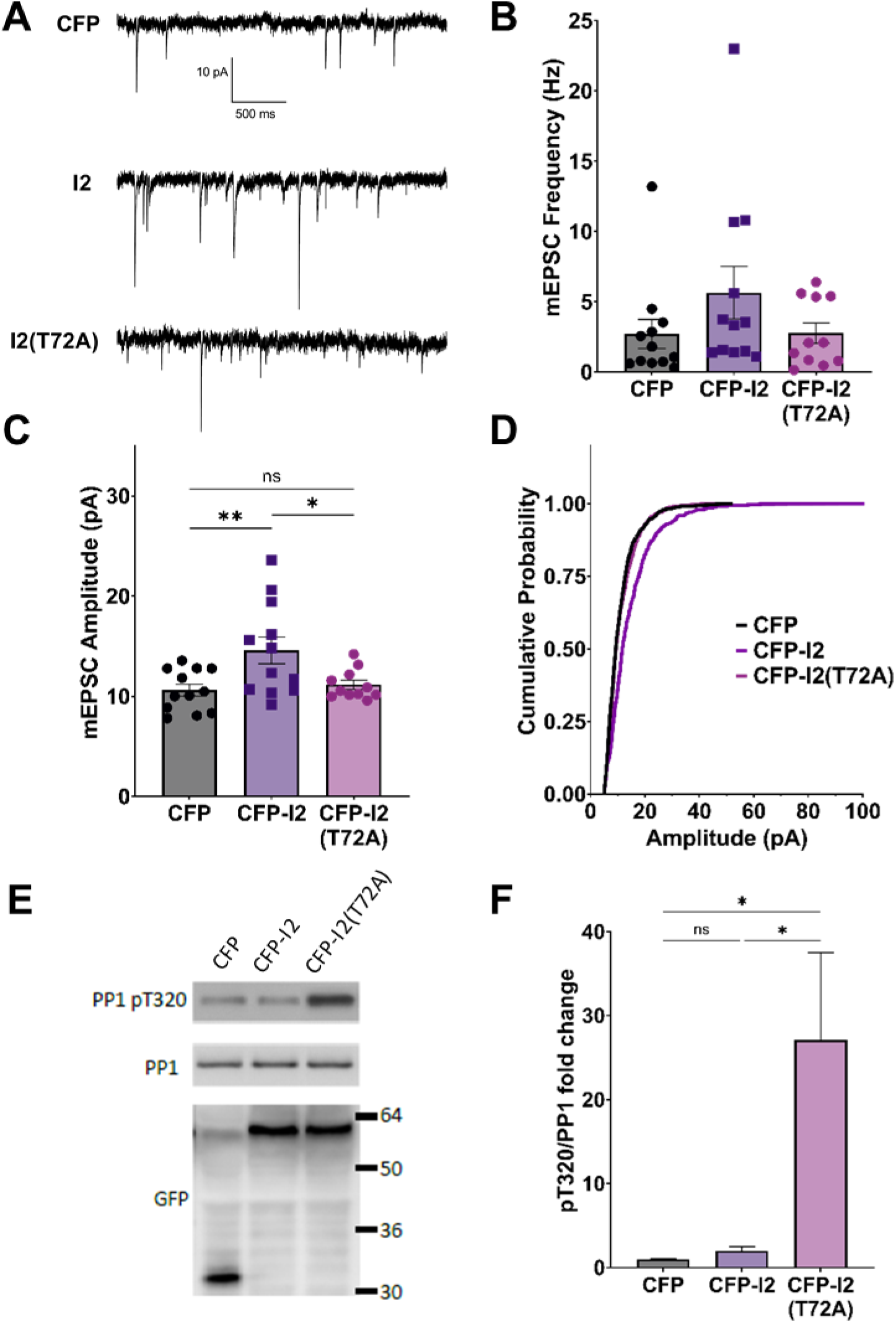
I-2 promotes synaptic transmission in primary cortical neurons. **A**: Example traces of mEPSC recordings in primary cortical neurons (∼DIV21) expressing CFP, CFP-I2, or CFP-I2(T72A). **B**,**C:** Quantification of mEPSC amplitude (B) and frequency (C). I-2 overexpression significantly increases mEPSC amplitude, but not frequency (one-way ANOVAs, F(2,32)=5.81, p<0.01; F(2,32)=1.62, p=0.21, respectively). **D:** Cumulative probability of mEPSC amplitude distribution (Kruskal Wallis statistic = 122.9, p<0.0001). Data in B-D are from the following number of cells, CFP: 12; I2: 12; I2(T72A): 11. **E:** Western blot derived from ∼DIV21 primary cortical neurons 24 hours after infection with CFP-, CFP-I2-, or CFP-I2(T72A)-expressing Sindbis virus. **F:** Quantification of western blot results from five sets of neuronal cultures. pT320 was first normalized to total PP1, then normalized to CFP control culture (one-way ANOVA, F(2,12)=6.01, p<0.05). Tukey post-hoc comparisons following one-way ANOVAs are displayed: *p<0.05, **p<0.01, ^ns^not significant.

In order to determine the effect of overexpressed I-2^WT^ and I-2^T72A^ on PP1 activity, we examined the status of endogenous PP1’s inhibitory phosphorylation at threonine 320 (pT320) [11]. Consistent with I-2^T72A^ as a constitutive PP1 inhibitor, we found that overexpressed CFP-I-2^T72A^ robustly (∼30 fold) and significantly increased pT320 (**Fig. 1E, F**), an inverse marker of PP1 activity [24]. Surprisingly, there was no significant difference in pT320 between CFP-I-2^WT^ and CFP (one-way ANOVA, Tukey post-hoc, n=5 per group, adjusted p=0.9927) (**Fig. 1E, F**), a drastic difference from the effect of CFP-I-2^T72A^ on pT320. This result is contrary to the simple interpretation of I-2 as a PP1 inhibitor from in vitro studies, but consistent with our previous report of robust PP1 activity in I-2 immunoprecipitates [13]. This is also consistent with the model of PP1 inhibition-activation by I-2: the rate-limiting step in the PP1 activation-inhibition cycle is the inhibition step [2] and the robust I-2 T72 kinase activity seen in neurons [11] should favor the activation state. On the other hand, I-2^T72A^ blocks the activation step and potently inhibits PP1 (**Fig. 1E, F**).

Since I-2^WT^ (**Fig. 1**) and the neurabin-PP1 holoenzyme [19] both promote basal synaptic transmission in neuronal cultures, we next sought to directly compare the function of I-2 and PP1 on synaptic transmission in acute hippocampal slices via genetic ablation. As neurabin has minimal binding to PP1β, and PP1β is not enriched in dendritic spines [25], we focused our study on PP1α and PP1γ. While mice with PP1α or PP1γ KO in neural progenitor cells (NPCs; Nestin-Cre, JAX) were viable, I-2 NPC KO was embryonic lethal. We therefore generated I-2 conditional KO mice with a hippocampus-specific Cre (CaMKII-cre^T-29^, JAX). We then measured basal synaptic transmission via input-output (I/O) curves at hippocampal CA3-CA1 Schaffer-collateral synapses in each transgenic mouse line. We found that synaptic transmission was significantly decreased in both I-2 and PP1γ KO mice compared to control littermates (**Fig. 2B, C; Fig. S1**). On the other hand, the I/O curve from PP1α KO mice was not significantly different compared to control littermates (**Fig. S1**), consistent with its lower binding affinity to neurabin. We did not observe a difference in paired-pulse facilitation (PPF) in any mouse model (**Fig. 2D, E; Fig. S1)**, indicating no change in glutamate release by CA3 neurons. These results thus suggest that I-2 and PP1γ, but not PP1α, promote AMPAR-mediated basal synaptic transmission in CA1 pyramidal neurons.

**Figure 2:**
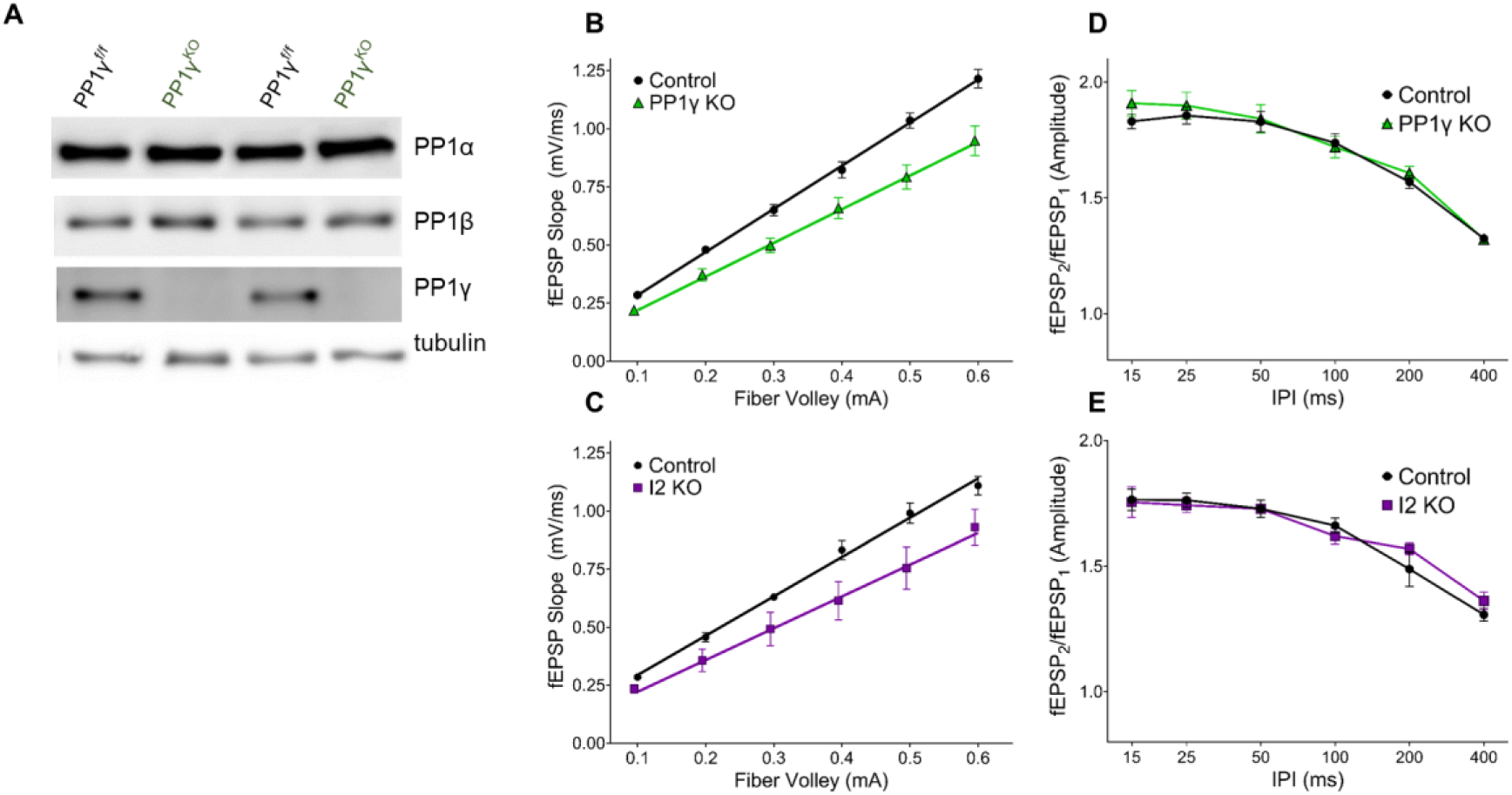
I-2 and PP1γ promote synaptic transmission in acute hippocampal slices. **A:** Successful knockout of PP1γ protein in nestin-cre;PP1γ^f/f^ mice. There is a slight upregulation of PP1β protein. **B-E**: Results from field recordings in acute hippocampal slices at Sch-CA1 synapses. **B**,**C**: There is a decrease in basal synaptic transmission in PP1γ^KO^ and I-2^KO^ mice (two-way RM-ANOVAs, genotype: F(1,16)=14.39, p<0.01; F(1,8)=6.987, p<0.05, respectively). **D**,**E:** There is no change in paired pulse facilitation (PPF) in PP1γ^KO^ or I-2^KO^ mice (two-way RM-ANOVAs, genotype: F(1,12)=0.280, p=0.606; F(1,8)=0.068, p=0.801, respectively). Data are from the following number of mice/slices: B: control 3/9, knockout 3/9; C: control 3/6, knockout 2/4; D: control 3/7, knockout 3/7; E: control 3/6, knockout 2/4.

Our data so far suggest that PP1γ and I-2^WT^, but not I-2^T72A^, promote excitatory synaptic transmission. Since we did not observe a change in PP1 activity by overexpressed I-2^WT^ (**Fig. 1E**), we next sought to determine the effect of I-2 on PP1 interaction with its regulatory proteins. We first established direct I-2– PP1 interaction via fluorescence lifetime imaging microscopy (FLIM). The fluorescence lifetime of CFP (Ƭ_CFP_) on CFP-I-2 was robustly decreased when YFP-PP1γ was co-expressed in HEK 293 cells (**Fig. 3A1-3**). Moreover, we found that the decrease of Ƭ_CFP-I-2_ was attenuated if YFP-PP1γ^H125Q^ was co-expressed, a PP1 mutant made to disrupt the binding interface between I-2 and PP1 [4] (**Fig. 3A**). This suggests that the FRET observed between CFP-I-2 and YFP-PP1γ is derived from I-2–PP1 interaction. We next examined the interaction between neurabin and I-2. We observed a robust decrease of Ƭ_CFP_ of CFP-neurabin^1-490^ when YFP-I-2 was co-expressed (**Fig. 3B1-3**). This decrease was not observed with a PP1-binding-deficient neurabin, CFP-neurabin^1-490, F460A^ (**Fig. 3B1-3**). Because the RvXF motif on neurabin does not participate in interaction with I-2 [15], this result suggests that the FRET between CFP-neurabin^1-490^ and YFP-I-2 involves a trimeric complex with endogenous PP1. We did not observe FRET between CFP-I-2 and neurabin^1-490^-YFP (**Fig. 3C1-3**), demonstrating that FRET between fluorescently-tagged I-2 and neurabin is sensitive to N- vs. C-terminal positioning of the fluorophores. However, we observed significant FRET between CFP-I-2^T72A^ and neurabin^1-490^-YFP (**Fig. 3C1-3**), supporting the idea that T72A mutation leads to a conformational change in I-2, large enough to position the fluorophores close enough and/or in optimal orientation for FRET to occur.

**Figure 3:**
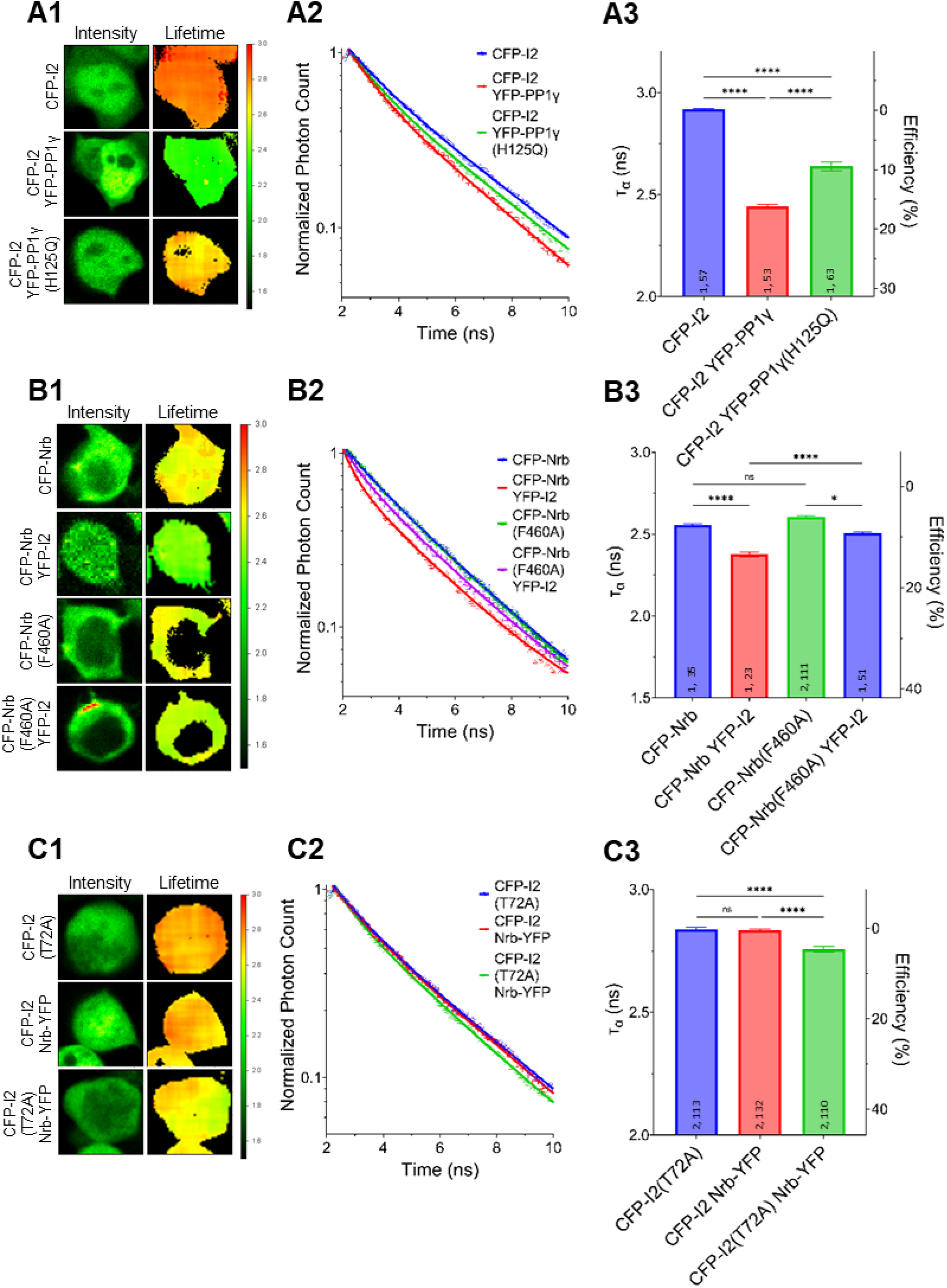
Regulation of I-2 interactions with PP1γ and neurabin in vivo as assessed by FLIM. **A1, B1, C1:** Intensity (left) and lifetime (right) of example cells. Lifetime expressed in nanoseconds (ns), with a color scale shown on the right. **A2, B2, C2:** Time-correlated single photon counting (TCSPC) histograms from example fields of view. **A3, B3, C3:** Mean lifetimes and SEM from the indicated constructs. **A3:** YFP-PP1γ, but not YFP-PP1γ^H125Q^, co-expression decreases CFP-I2 lifetime (one-way ANOVA, F(4,289)=417.9, p<0.0001). **B3:** YFP-I2 co-expression decreases CFP-Nrb lifetime. The interaction between CFP-Nrb^1-490^and YFP-I2 is attenuated by PP1-binding-deficient Nrb, CFP-Nrb^1-490, F60A^ (one-way ANOVA, F(3,216)=35.60, p<0.0001). **C3:** Nrb^1-490^-YFP does not affect CFP-I2 lifetime, but decreases CFP-I2^T72A^ lifetime (one-way ANOVA, F(2,352)=27.86, p<0.0001). Tukey post-hoc comparisons following one-way ANOVAs are displayed: *p<0.05, **p<0.01, ***p<0.001, ****p<0.0001, ^ns^not significant. Number of cultures and number of cells are shown within bars. All data represent independent experiments; lifetime data was not reused between subfigures.

We next determined whether I-2 can promote neurabin-PP1γ holoenzyme formation. We observed a robust decrease of Ƭ_CFP_ on CFP-neurabin^1-490^ when it was co-expressed with YFP-PP1γ (**Fig. 4A**). Moreover, the decrease of Ƭ_CFP_ was significantly reduced when YFP-PP1γ was co-expressed with PP1-binding deficient CFP-neurabin^1-490, F460A^ (**Fig. 4A**). This suggests that the FRET observed between CFP-neurabin^1-490^ and YFP-PP1γ is derived from their interaction. Notably, we found that co-expressing rLuc-I-2^WT^ with CFP-neurabin (1-490) and YFP-PP1γ significantly further decreased the Ƭ_CFP_ on CFP-neurabin^1-490^, relative to CFP-neurabin^1-490^ and YFP-PP1γ double transfection (**Fig. 4B**). This suggests that I-2 promotes neurabin-PP1γ holoenzyme formation, thus regulating PP1γ function in a positive manner. This finding is consistent with previous in vitro binding data showing I-2 promotes neurabin-PP1 binding [16]. A similar study of I-2 function in tobacco leaves showed that I-2 promotes PP1 binding to its scaffolding protein SNF1-related protein kinase 2 (SnRK2) [26]. Overall, our study suggests that overexpressed I-2^WT^ increases synaptic transmission, not via promoting the enzymatic activity of individual PP1 molecules (**Fig. 1D, E**), rather via increasing the number of PP1 molecules targeted by neurabin.

**Figure 4:**
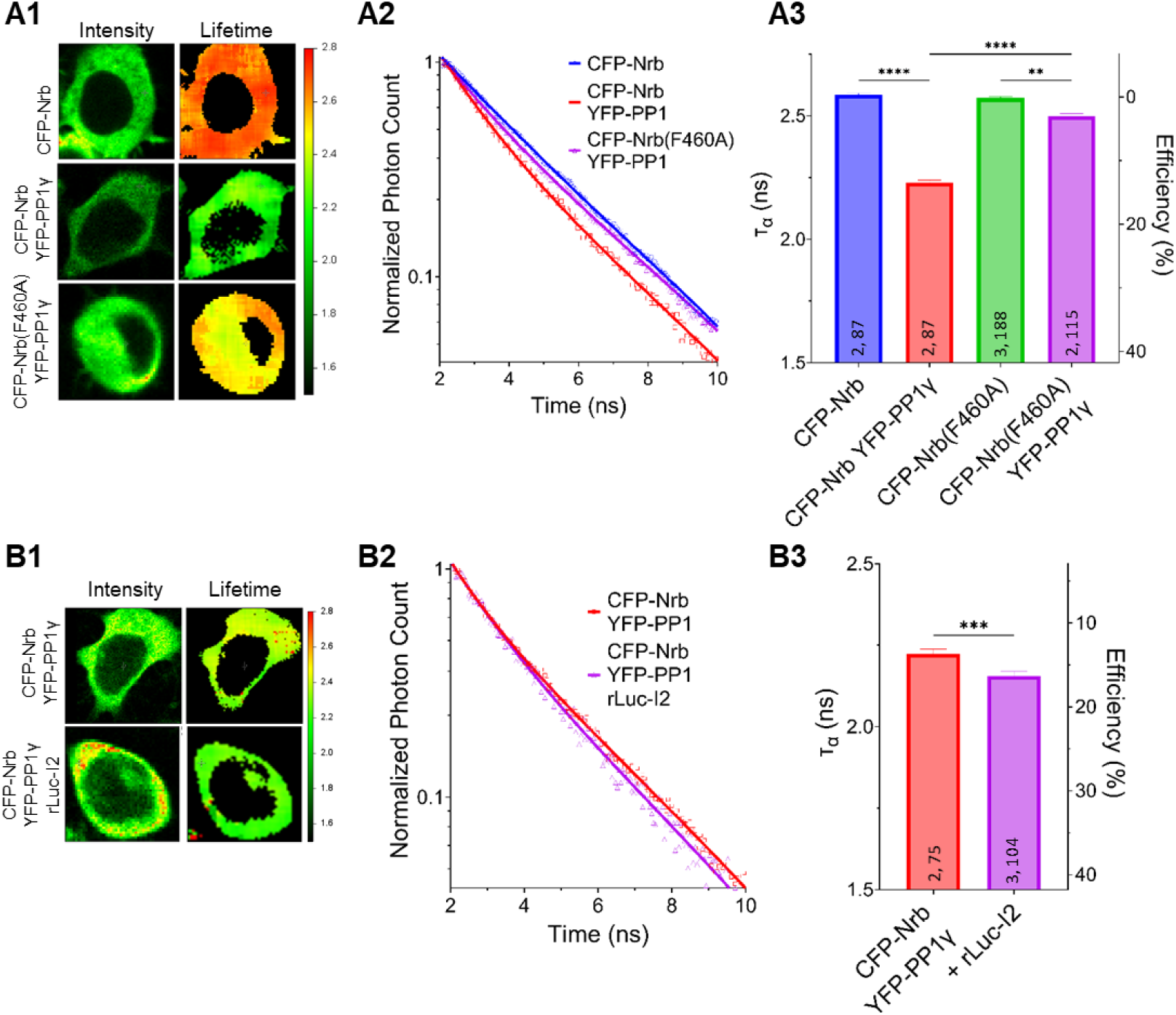
I-2 promotes neurabin-PP1 interaction. **(A1, B1)** Intensity (left) and lifetime (right) of example cells. Lifetime expressed in ns. **(A2, B2)** Time-correlated single photon counting (TCSPC) histograms from example fields of view. **(A3, B3)** Mean lifetimes and SEM from the indicated constructs. **A3:** YFP-PP1γ co-expression decreases CFP-Nrb^1-490^ lifetime. CFP-Nrb^1-490, F460A^ significantly attenuates the YFP-PP1γ-mediated decrease in lifetime (one-way ANOVA, F(7,694)=190.4, p<0.0001). **B3:** rLuc-I2 co-expression promotes CFP-Nrb^1-490^ interaction with YFP-PP1γ, further decreasing lifetime (two-tailed t-test, *t*(177)=3.443, p<0.001). t-test results (B3) or Tukey post-hoc comparisons following one-way ANOVAs (A3) are displayed: *p<0.05, **p<0.01, ***p<0.001, ****p<0.0001, ^ns^not significant. Number of cultures and number of cells are shown within bars. All data represent independent experiments; lifetime data was not reused between subfigures.

## Conclusions

In summary, we found that I-2 and PP1γ, but not PP1α, promotes basal synaptic transmission and that I-2 positively regulates PP1γ function via enhancing neurabin-PP1γ holoenzyme formation, without substantially affecting PP1γ enzymatic activity. Our study provides a fundamental mechanism by which I-2 can positively regulate PP1 function. This mechanism may regulate in vivo PP1 function beyond synaptic transmission.

## Materials & Methods

### Conditional knockout mice

Nestin-cre and CaMKIIα-cre (T29-1) were purchased from Jackson Lab. PP1α and PP1γ conditional KO mouse were generated as described previously [27]. I2 floxed mice, in which exon 1 and exon 2 were flanked by Cre recombinase-dependent loxP recognition sequences, were generated by the University of Rochester Medical Center Transgenic and Genome Editing Core.

### Primary neuronal cell cultures, infections, immunoblotting and antibodies

Primary cortical neurons were prepared from mixed male/female E18 Sprague Dawley (SD) rat embryos as previously described [28, 29]. ∼DIV21 neurons were used in our study. CFP, CFP-I-2(WT) and CFP-I-2(T72A) constructs and recombinant Sindbis virus generation and infection have been described previously [11]. In brief, pSinRep5 (nsP2S) vector was used for CFP/I-2 subcloning, and targeted recombinant constructs were linearized, in vitro translated and electroplated into BHK21 cells, along with helper DHBB. Supernatant containing recombinant viruses were collected, concentrated via centrifugation before being applied to cultured neurons for 24 hours. Medium from the 12-well neuronal plates were aspirated quickly, before 1X SDS gel loading buffer (contains protease and phosphatase inhibitor) was added to each well for about 10 min on ice before the cell lysates were harvested and heated for 10 min at 95°C followed by SDS-PAGE and immunoblotting. Anti-PP1 pT320 (1:1,000; Cell Signaling Technology), anti-PP1 (1:1,000; E-9, Santa Cruz Biotechnology, Inc.), and anti-GFP (1:1000, 0.4 µg/ml; Roche).

### Electrophysiology

Whole-cell patch-clamp recordings on cultured cortical neurons were recorded at ∼DIV21, as described [14]. Neurons were transfected with CFP, CFP-I-2^WT^ or CFP-I-2^T72A^ by calcium phosphate precipitation method 3 days prior. Pipettes were filled with (in mM): 117 Cs-methylsulfonate, 20 HEPES, 1 EGTA, 0.1 CaCl2, 5 CsCl, 2.5 MgATP, 0.25 Na3GTP, pH 7.4. External solution consisted of (in mM): 135 NaCl, 3.5 KCl, 2 CaCl2, 1.3 MgCl2, 10 HEPES, 20 glucose, supplemented with 200 nM TTX, 25 µM D-AP5, and 50 µM picrotoxin. mEPSCs were detected using template fitting in Clampfit 10.3 with a 5 pA threshold. Cumulative distributions were generated by histogram cumulative distribution of all mEPSC events for CFP (981 events) and I2^T72A^ (932 events) groups. An equivalent number of events was randomly sampled from each I-2^WT^ cell, yielding 946 events.

Acute hippocampal slices were prepared from 1-2-month old mice bred as previously described [30]. 400 μm thick slices were prepared after decapitation and rapid extraction of the brains into ice-cold artificial cerebrospinal fluid (ACSF). Slices were then recovered in room temperature (RT) ACSF for at least 1 hr prior to field recordings. Recordings were conducted at Schaffer collateral-CA1 synapses in RT ACSF at a flow rate of 2-3 mL/min. A borosilicate recording electrode (1-3 MΩ) filled with 1 M NaCl was placed in CA1 stratum radiatum and a monopolar borosilicate filled with ACSF (Fig. 2) or tungsten concentric bipolar stimulating electrode (FHC) (Fig. S2) placed on Schaffer collaterals between CA3 and CA1. Responses were elicited every 15 sec, with stimulation delivered by an ISO-Flex stimulus isolator (AMPI). The ACSF solution consisted of (in mM): 126.0 NaCl; 2.5 KCl; 2.5 CaCl_2_; 1.3 MgSO_4_; 1.25 NaH_2_PO_4_; 26.0 NaHCO_3_; and 10.0 D-glucose. ACSF was continuously aerated with carbogen (95% O2, 5% CO2) during incubation and recordings. Basal synaptic transmission was assessed by input-output (IO) curves. Short-term pre-synaptic plasticity was assayed using paired-pulse facilitation (PPF) of varying inter-pulse intervals. Recordings were collected with a MultiClamp 700A amplifier (Axon Instruments), PCI-6221 data acquisition device (National Instruments), and Igor Pro 7 (Wavemetrics) with a customized software package (Recording Artist, http://github.com/rgerkin/recording-artist).

### Fluorescence Lifetime Imaging Microscopy (FLIM)

HEK293 cells were transfected in 35 mm culture dishes and imaging was performed 2 days later. Images were captured on an Olympus BX51WI upright microscope using a water-immersion 25x objective (Olympus XLPlan N). Two-photon 850 nm excitation was achieved with a Mai Tai Ti:Sapphire multi-photon laser (Spectra Physics), using a repetition rate of 80 MHz and a pulse width of approximately 100 fs, and emission was filtered with a 480-20 filter. Emission was measured by a H72422P Hamamatsu hybrid avalanche photodiode. Time-correlated single photon counting was performed using a Becker and Hickl module with a 25 ps resolution. VistaVision software (ISS) was used for lifetime analysis. Donor fluorescence from individual cells was binned and fit with a double exponential function, consistent with the lifetime properties of CFP.

### Analysis

GraphPad Prism (9.3.0) was used for statistical analyses and data visualization. Statistical significance between means was calculated using unpaired, two-tailed t-tests or ANOVAs. Repeated-measure (RM)-ANOVAs were used for IO and PPF comparisons. Tukey and Bonferroni post-hoc comparisons were performed for one- and two-way ANOVAs, respectively. The arithmetic mean and standard error of the mean are displayed in all figures unless otherwise specified.

## Acknowledgments

This work was supported by the National Institutes of Health (NIH) R01 MH109719 and National Science Foundation (NSF) IOS-1457336 to H.X.; NIH DA10044 to A.C.N.; NIH F30 MH122046 to K.F.; NIH R35GM137951 to D.M.M.; the Heart and Stroke Foundation, Canada, to H.A; and the Natural Sciences and Engineering Research Council of Canada to D.S. The authors declare no competing financial interests.

## Figure legend

**Figure S1:**
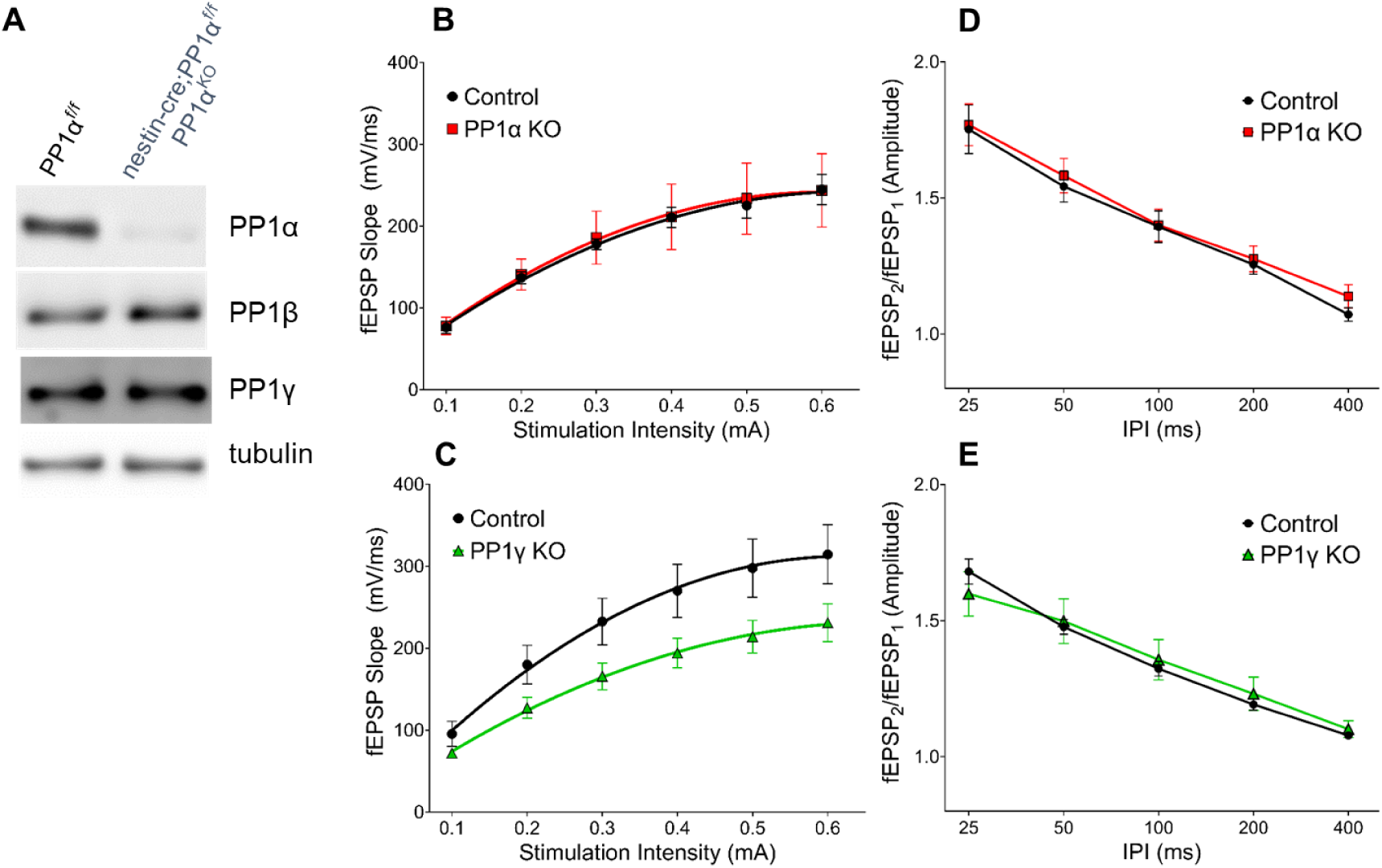
PP1α does not play a role in synaptic transmission. **A:** Successful knockout of PP1α protein in nestin-cre;PP1α^f/f^ mice. **B-E**: Results from field recordings in acute hippocampal slices at Sch-CA1 synapses. **B**,**C:** There is no difference in basal synaptic transmission in PP1α^KO^ mice, but a significant decrease in PP1γ^KO^ mice (two-way RM-ANOVAs, genotype: F(1,10)=0.02, p=0.899; F(1,14)=4.79, p<0.05, respectively). **D**,**E:** There is no change in paired pulse facilitation (PPF) at the Sch-CA1 pyramidal synapses in PP1α^KO^ or PP1γ^KO^ mice (two-way RM-ANOVAs, genotype: F(1,10)=0.172, p=0.687; F(1,14)=0.006, p=0.938, respectively). Data are from the following number of mice/slices: B: control 2/7, knockout 2/5; C: control 3/7, knockout 3/9; D: control 2/6, knockout 2/6; E: control 3/6, knockout 3/10.

